# Accurate cytogenetic biodosimetry through automation of dicentric chromosome curation and metaphase cell selection

**DOI:** 10.1101/120410

**Authors:** Jin Liu, Yanxin Li, Ruth Wilkins, Farrah Flegal, Joan H. Knoll, Peter K. Rogan

**Affiliations:** Department of Biochemistry, University of Western Ontario, Health Canada; Cytognomix Inc, Health Canada.; Radiobiology and Protection Division, Health Canada; Canadian Nuclear Laboratories; Department of Pathology and Laboratory Medicine, University of Western Ontario

**Author notes:** Correspondence: Peter K. Rogan, Ph.D. Department of Biochemistry Schulich School of Medicine and Dentistry University of Western Ontario London, Ontario N6A 5C1 Canada E. **ABBREVIATIONS** ADCI: *Automated Dicentric Chromosome Identifier* CNL: *Canadian Nuclear Laboratories* DC: *Dicentric chromosome* DCA: *Dicentric chromosome assay* FP: *False positive dicentric chromosome* HC: *Health Canada* K–S: *Kolmogorov–Smirnov test* MC: *Monocentric chromosome* MC-DC SVM: *Monocentric-Dicentric Support Vector Machine* ML: *Machine learning* SCS: *Sister chromatid separation* SD: *Standard deviation* SVM: *Support Vector Machine* TP: *True positive dicentric chromosome.

## Abstract

Software to automate digital pathology relies on image quality and the rates of false positive and negative objects in these images. Cytogenetic biodosimetry detects dicentric chromosomes (DCs) that arise from exposure to ionizing radiation, and determines radiation dose received from the frequency of DCs. We present image segmentation methods to rank high quality cytogenetic images and eliminate suboptimal metaphase cell data based on novel quality measures. Improvements in DC recognition increase the accuracy of dose estimates, by reducing false positive (FP) DC detection. A set of chromosome morphology segmentation methods selectively filtered out false DCs, arising primarily from extended prometaphase chromosomes, sister chromatid separation and chromosome fragmentation. This reduced FPs by 55% and was highly specific to the abnormal structures (≥97.7%). Additional procedures were then developed to fully automate image review, resulting in 6 image-level filters that, when combined, selectively remove images with consistently unparsable or incorrectly segmented chromosome morphologies. Overall, these filters can eliminate half of the FPs detected by manual image review. Optimal image selection and FP DCs are minimized by combining multiple feature based segmentation filters and a novel image sorting procedure based on the known distribution of chromosome lengths. Applying the same segmentation filtering procedures to both calibration and test sample image data reduced the average dose estimation error from 0.4Gy to <0.2Gy, obviating the need to first manually review these images. This reliable and scalable solution enables batch processing multiple samples of unknown dose, and meets current requirements for triage radiation biodosimetry of high quality metaphase cell preparations.

## INTRODUCTION

The analysis of microscopy images of cells is the basis of several types of analysis of the effects of damage by ionizing radiation. The gold standard radiation biodosimetry method, the dicentric chromosome assay (DCA), involves measuring the frequency of aberrant dicentric chromosomes in a patient sample. While some aspects of the assay have been successfully automated and streamlined, its overall throughput remains limited by the labour-intensive dicentric (DC) scoring step, potentially affecting timely estimation of radiation exposures of multiple affected individuals, for example, in a large accident or a mass casualty event^1,2^.

One issue with automated analysis is the selection of images of adequate quality for accurate identification of the chromosome damage. With DCA, the decision to select or exclude microscope images for analysis has traditionally been performed manually; yet current automated image capture approaches make this approach impractical due to the growing size of datasets. Image quality assessment often estimates new data in relation to reference images^3^, complex mathematical models^4^, or distortions from a training set recognized by machine learning^5^. Generic methods of assessing image quality are not appropriate in our situation. Features tailored for ranking chromosome images cannot be generalized to entropy measures based on applying frequency filter to intensity distributions. To be useful, quality assurance for evaluation of specific microscopic biological objects in an image may require expert-derived rules to categorize preferred images.

To address issues with automation of the DCA, we have been developing the Automated Dicentric Chromosome Identifier (ADCI) software to automate DC scoring and radiation dose estimation. The algorithms underlying ADCI have been described and experimentally validated^6–11^. Briefly, foreground objects are extracted from the metaphase cell image by thresholding intensities above background levels. Preprocessing filters remove most (but not all) non-chromosomal objects (e.g. debris, nuclei, overlapping chromosomes). Each remaining object is regarded as a single, intact, post-replication “chromosome” object. Each chromosome is processed to determine a contour (chromosome boundary) and its centerline (chromosome long axis). The Intensity-Integrated Laplacian method^9,10^ constructs a width profile from consecutive vector field tracelines running approximately orthogonal to the centerline, and potential centromere locations (“centromere candidates”) are identified from constrictions in the said width profile (see Fig. 1)^12^. Machine learning (ML) modules use image segmentation features derived from each chromosome to classify centromeres and dicentric chromosomes^6,11^. The first Support Vector Machine (SVM) ranks potential centromere candidates in each chromosome according to their corresponding hyperplane distances; then another SVM scores the chromosome as either monocentric (MC) or dicentric (DC) using features derived from the top two candidates.

**Figure 1.**
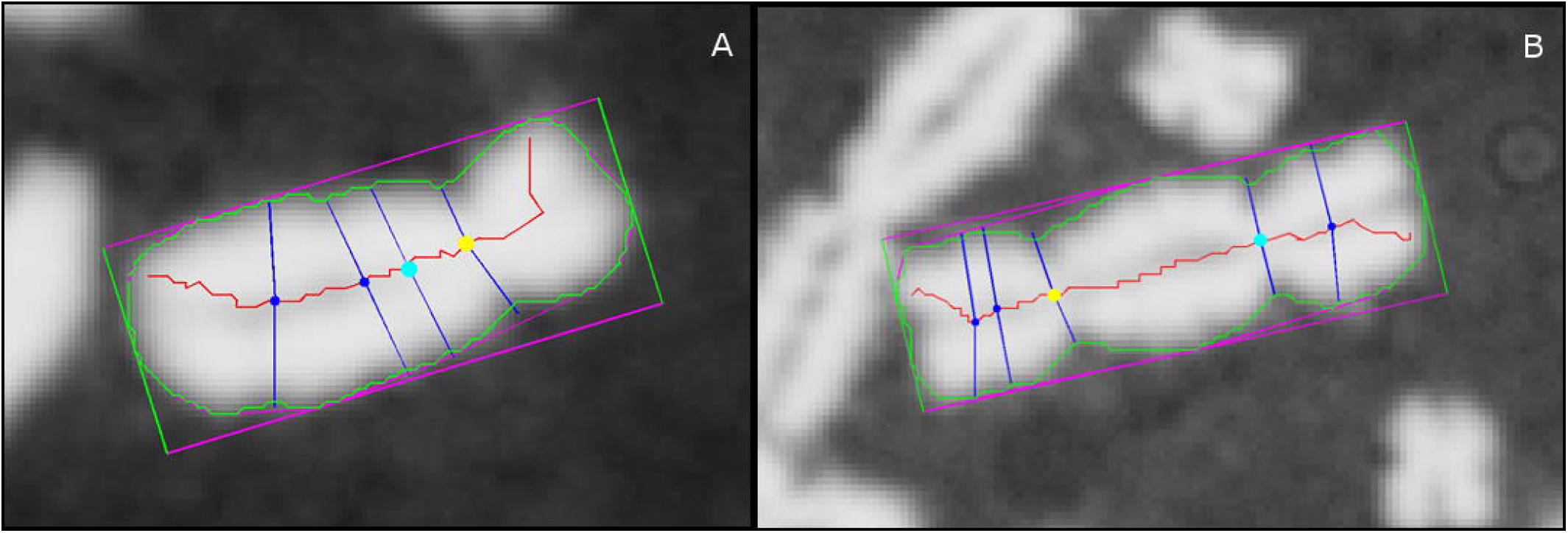
Chromosome images processed by ADCI, annotated with key segmentation features. (A) Monocentric and (B) Dicentric chromosome. Chromosome contour is overlaid in green, long-axis centreline in red. Yellow and cyan markers on the centerline indicate the top-ranked and 2^nd^-ranked centromere candidate, respectively (other candidates not shown), with their corresponding width tracelines (roughly orthogonal to centerline) displayed in the same colour. Arc lengths of width tracelines running down the centerline (not all shown) are used to construct a chromosomal width profile. Note that the top-ranked candidate correctly labels the true centromere location, while the 2^nd^-ranked candidate labels a minor non-centromeric constriction. By comparing features extracted from both candidates (including width and pixel intensity information), the software correctly assessed that only one of the candidates is an actual centromere, so the chromosome was classified as monocentric. In dicentric chromosomes, both candidates would label actual centromeres.

Samples exposed to known radiation doses (in Gy) are processed by ADCI to construct a dose-response calibration curve. The average frequency of DC’s per cell in dose calibrated samples, the radiation response, is fit to a linear-quadratic function. The response for test samples exposed to unknown radiation levels can then be analyzed with this equation to estimate their corresponding doses.

We noticed that metaphase cell images of inconsistent quality can affect accuracy of dose estimation by ADCI. Previous studies evaluated the efficacy of ADCI at chromosome classification and dose estimation^10,11^. While the sensitivity (recall) for DCs was acceptable (∼70%) and relatively constant at all radiation exposure levels, precision showed a strong dependence on dose. Chromosome misclassification, in particular false positive dicentrics (FPs) were more prevalent at low (≤1Gy) compared to high (3–4Gy) doses; at 1Gy, FPs could outnumber true positive dicentrics (TPs) by a factor of 4 to 5. Consequently, ADCI-processed samples exhibited a reduced range of accurate responses to radiation compared to manually scored samples. Although use of the same algorithm to derive the calibration curve compensates for some of these differences, reliability of dose estimation ultimately hinges on DC classification accuracy. As DCs are greatly outnumbered by MCs (background frequency in normal, unexposed individuals is one DC per 1000 cells^6^), this study focuses on improving the distinction between TP and FP DCs without compromising recall.

FPs reflect inadequacies in misinterpreting certain chromosome morphologies or non-chromosomal objects. Selective targeting and removal of these instances would reduce FPs without limiting TP identification, improving overall classification accuracy. We investigated FP morphologies to identify problematic cases, and devised a set of post-processing object segmentation filters to eliminate them. Then, to ensure consistent performance, segmentation filters were developed to remove poor quality cell images. These images are usually characterized by either a lack of or incomplete complement of metaphase chromosomes, misclassified interphase or micro-nuclei as metaphases, or incorrectly segmented sister chromatids as individual chromosomes. Each proposed filter was tested individually, and the best performing filters were integrated, and tested on actual cytogenetic dosimetry data exposed to various radiation doses. The effects of these filters on classification performance was evaluated on image sets from two independent biodosimetry laboratories, and their impact on dose estimation was assessed on cells obtained from an international biodosimetry exercise.

We present this hybrid approach which selects images based on a combination of optimal global image properties for scoring metaphase cells, and customized object segmentation, identification and elimination of false positive DCs. These improvements in ADCI ensure timely, reproducible, and accurate quantitative assessment of acute radiation exposure.

## METHODS & MATERIALS

Cytogenetic data were obtained by biodosimetry laboratories at Health Canada (HC) and Canadian Nuclear Laboratories (CNL) according to IAEA guidelines. Blood samples were irradiated by an XRAD-320 (Precision X-ray, North Branford, CT) at Health Canada and processed at both laboratories. Peripheral blood lymphocyte samples were cultured, fixed, and stained at each facility according to established protocols^2,12^. Metaphase images from Giemsa-stained slides were captured independently by each lab using an automated microscopy system (Metasystems). One set of metaphase images from CNL and two sets from HC (Table 1) were used for development and initial testing of the proposed algorithms. After image processing by ADCI, called DCs were manually reviewed and the consensus scores of TPs or FPs by 3 trained individuals were determined. Calibration curves were prepared based on 6 samples of known radiation dose (Table 2). An additional 6 samples^11^ were initially blinded to the actual radiation exposures as test samples (Table 3). Test samples were exposed to a range of radiation doses bounded by the doses of samples used to construct the calibration curve. The sample naming convention is the lab name followed by the sample identifier, e.g. HC1Gy signifies the 1 Gy calibration sample prepared at HC, whereas CNL-INTC03S04 represents the INTC03S04 international exercise test sample (exposed to 1.8 Gy) prepared at CNL.

**Table 1.**
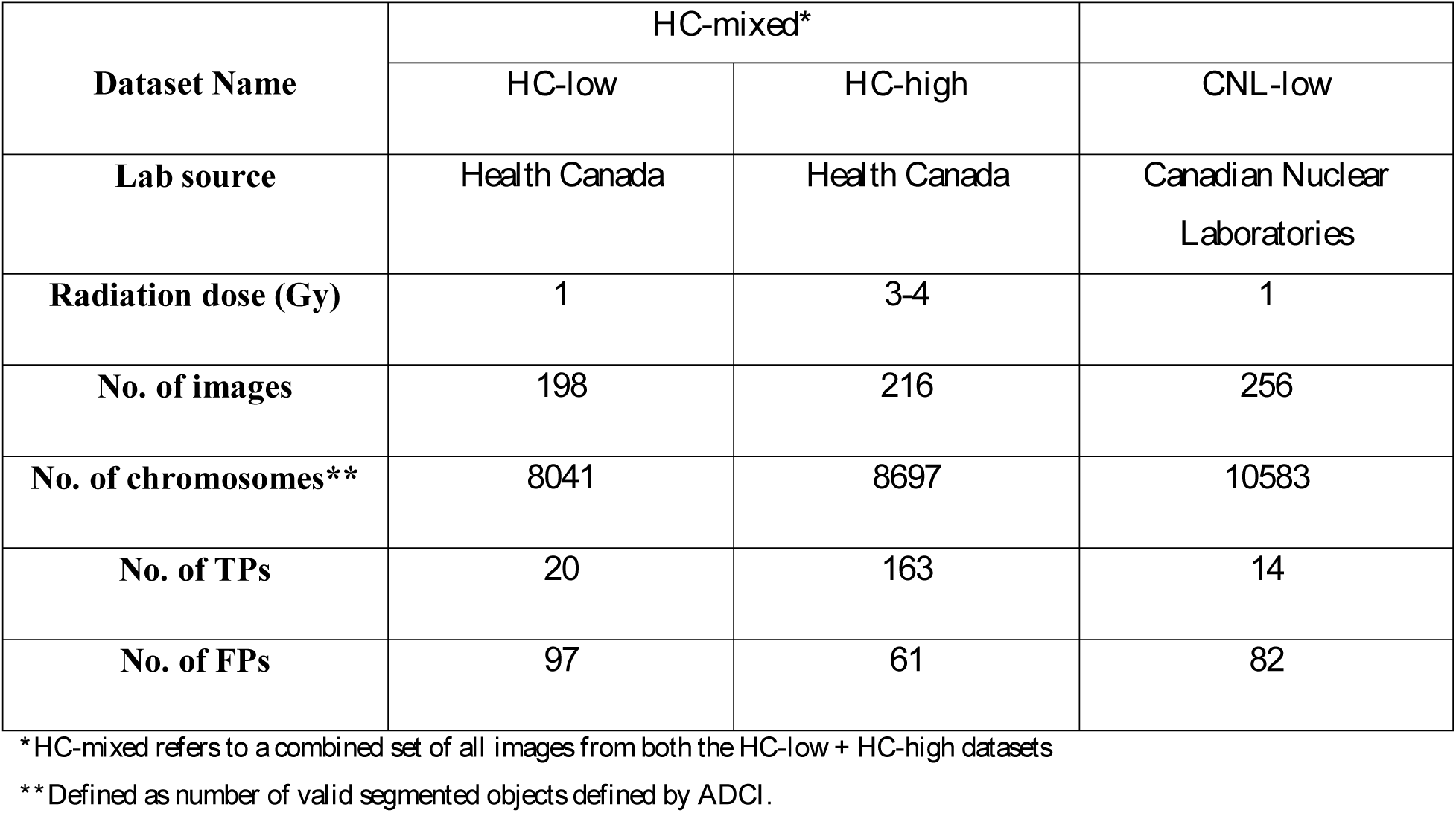
Metaphase image sets used in development and validation of DC filters.

| Dataset Name | *HC-mixed\** |  |  |
| --- | --- | --- | --- |
|  | *HC-low* | *HC-high* | *CNL-low* |
| **Lab source** | Health Canada | Health Canada | Canadian Nuclear Laboratories |
| **Radiation dose (Gy)** | 1 | 3-4 | 1 |
| **No. of images** | 198 | 216 | 256 |
| **No. of chromosomes\*\*** | 8041 | 8697 | 10583 |
| **No. of TPs** | 20 | 163 | 14 |
| **No. of FPs** | 97 | 61 | 82 |

\**HC-mixed* refers to a combined set of all images from both the *HC-low* + *HC-high* datasets
\*\*Defined as number of valid segmented objects defined by ADCI.

**Table 2.**
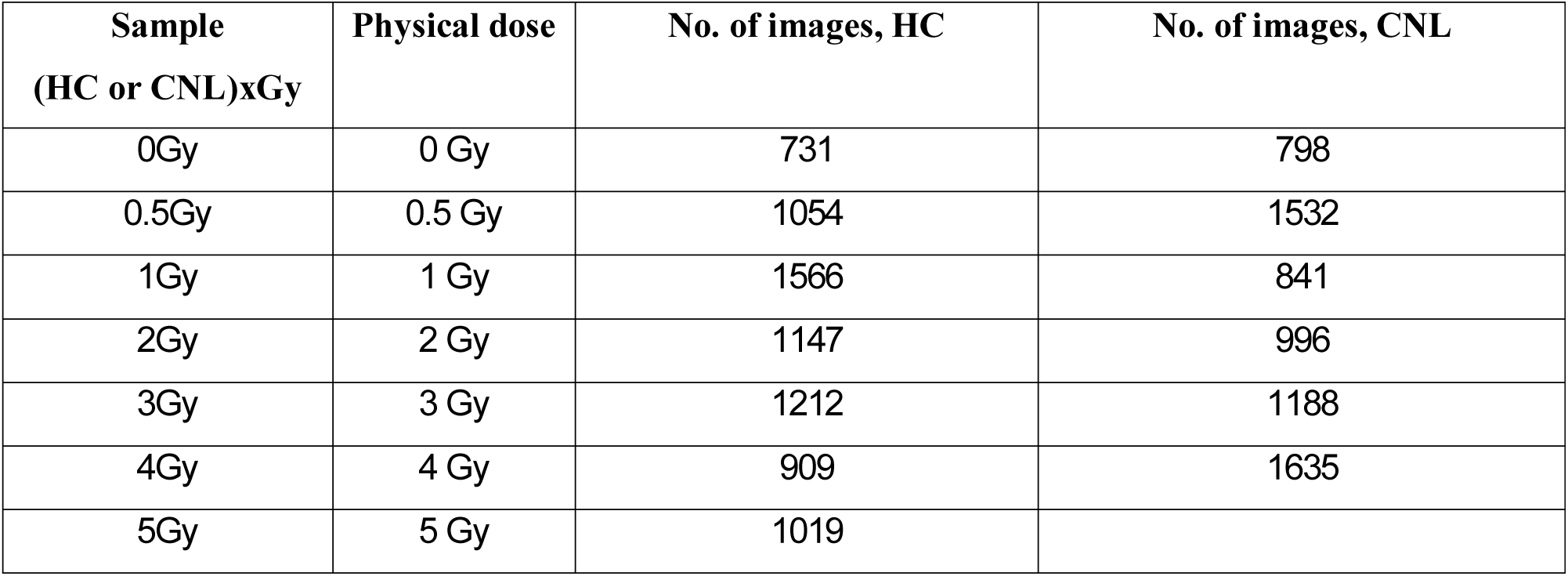
Metaphase image samples used in construction of dose calibration curves.

**Table 3.** Metaphase image samples used in evaluation of dose assessment performance.

| Sample name | Physical Dose<br>(Gy) | No. of images, HC<sup>1</sup><br>preparation | No. of images, CNL<sup>2</sup><br>preparation |
| --- | --- | --- | --- |
| INTC03S01 | 3.1 | 540 | 500 |
| INTC03S08 | 2.3 | 637 | 500 |
| INTC03S10 | 1.4 | 708 | n/a |
| INTC03S04 | 1.8 | 600 | 957 |
| INTC03S05 | 2.8 | 1136 | 1527 |
| INTC03S07 | 3.4 | 477 | 735 |

<sup>1</sup>HC: Health Canada. <sup>2</sup>CNL: Canadian Nuclear Laboratory. n/a: sample data were not available.

Data consisted either of all “metaphase” images captured by the microscopy system, or a manually curated set of 500 high quality images. Selection of raw metaphase images for inclusion in samples was done automatically at HC using the default image classifier of the Metafer slide scanning system, while CNL selected images manually according to IAEA guidelines^12^. Experts from CNL selected for images deemed analyzable by humans with respect to chromosome count, spatial distribution and morphology.

### 1) ADCI settings & metaphase image data

ADCI software (V1.0)^11^ was used for DC detection and dose prediction, with the MC-DC SVM tuning parameter, σ, set to 1.5. ADCI libraries were initially written in MATLAB (R2014a) to develop and test the proposed DC FP filters, and were subsequently rewritten in C++ and integrated into ADCI. For development and validation of segmentation filters, independent datasets used three sets of roughly 200 images each (2 low dose, 1 high dose) were prepared from larger image sets that were originally used for validation of previous versions of ADCI (see Table 1; *HC-mixed* image set).

### 2) Morphological characterization of FPs

FPs and TPs were compared according to their respective segmentation features, including contour, width profile, centerline placement, centromere candidate placement, and total pixel area (Table 1). FPs were grouped by common distinguishing traits and assigned to one or more of the following morphological classes:

I. **Sister chromatid separation:** Sister chromatid separation (SCS) of a chromosome refers to the loss of sister chromatid cohesion at the telomeres, and often along the sister chromatids, excluding the centromeres. Due to inherent limitations of a centerline derived from contour skeletonization in chromosomes, SCS often resulted in partial or complete localization of the centerline along a single chromatid, rather than along the long axis of the full-width chromosome^8–10^. Complete centerline localization to chromatids of the q arm was common among acrocentric chromosomes (see Fig. 2A). This resulted in a width profile in which the displaced centerline did not accurately represent the width of the chromosome, and compromised centromere determination.
II. **Chromosome fragmentation:** Sister chromatid pairs were completely dissociated in metaphase images, resulting in incorrect labeling of each chromatid as separate chromosomes. Occasionally, segmentation fragmented images of intact non-uniform chromosomes into multiple, chromosomal artifacts^6^ (see Fig. 2B). Artifactual fragmentation into incomplete chromosome fragments led to unpredictable results, increasing FPs and FNs.
III. **Chromosome overlap:** Poor spatial separation of chromosomes produced clusters of overlapping/touching chromosome clusters which were inseparable. Occasionally, the cluster is segmented as a single contiguous object (see Fig. 2C). Like chromosome fragments, analysis of these overlapping chromosome clusters produces erroneous results. FP DCs were produced from clusters comprising two underlying monocentric chromosomes, each contributing a centromere to the combined object.
IV. **Noisy contour:** Poor image contrast at the chromosomal boundary produced “noisy,” jagged chromosome contours contributing multiple small constrictions to the width profile (see Fig. 2D). These artifactual constrictions were incorrectly identified as multiple centromeres if their magnitudes were similar to the true centromere, leading to FP assignment.
V. **Cellular debris:** Non-chromosomal objects such as nuclei and cellular debris were generally removed by pre-processing based on thresholding relative size and pixel intensity. However, aggregated cellular debris were occasionally labelled as a chromosome and naively analyzed by the software (see Fig. 2E).
VI. **Machine learning error:** A “catch-all” subclass for MCs with no identifiable morphological traits and reasonable contours and centerlines (see Fig. 2F). These cases reflect deficiencies in the feature set or training data of the machine learning (ML) classifiers, rather than image segmentation errors.

### 3) Filtering out False Positive Objects

Quantitative filters were created and tested to delineate FP DCs. Each formula targets one or more of the morphological classes described above, and generates a unitless filter score for each object, independent of the biodosimetry reference laboratory source. For any metaphase image, *{c*_*1*_*,…,c*_*N*_*}* denotes the set of N chromosomes within the image and *c\** denotes the predicted DC of interest. Each filter classifies c* as either a TP or FP by comparing its filter score against a heuristically-defined threshold that is independent of laboratory provenance. Thresholds were established empirically to maximize elimination of FPs without altering recognition of TPs. FPs generally produce lower filter scores than TPs (i.e. lower area, lower width, less oblong footprint, more asymmetrical), so FPs were selectively targeted by eliminating candidate DCs with scores below a threshold. Due to the low frequency of DCs in any given sample, minimizing the loss of TPs is paramount to minimize the likelihood of TP removal. For each filter, corresponding filter scores were calculated for all DCs in the *HC-mixed* image set (Table 1), and a heuristic threshold (to 2 significant digits; see below) was set to the minimum value observed in TPs. Thresholds for filters VI to VIII were calculated by repeating the same procedure on a chromosome set of 244 TPs from the MC-DC SVM training set, and the final thresholds were set to the lower of each pair of values.

I. **Area filter:** *A(c)* denotes the pixel area occupied by chromosome *c* (see Fig 3B). *c\** was classified as FP if *A(c*)/median({A(c*_*1*_*),…,A(c*_*N*_*)}) < 0.74* or as TP otherwise. This filter targets small acrocentric chromosomes (commonly displaying SCS) and chromosome fragments.
II. **Mean width filter:** *W*_*mean*_*(c)* denotes the mean value of the width profile of chromosome *c* (see Fig 3C). *c\** was classified as FP if *W*_*mean*_*(c*)/median({W*_*mean*_*(c*_*1*_*),…,W*_*mean*_*(c*_*N*_*)}) < 0.80* or as TP otherwise. This filter targets SCS and chromosome fragments.
III. **Median width filter:** *W*_*med*_*(c)* denote the median value of the width profile of chromosome *c* (see Fig 3C). *c\** was classified as FP if *W*_*med*_*(c*)/median({W*_*med*_*(c*_*1*_*),…,W*_*med*_*(c*_*N*_*)}) < 0.77* or as TP otherwise. This filter targets SCS and chromosome fragments.
IV. **Max width filter:** *W*_*max*_*(c)* denotes the maximum value of the width profile of chromosome *c* (see Fig. 3C). *c\** was classified as FP if *W*_*max*_*(c*)/median({W*_*max*_*(c*_*1*_*),…,W*_*max*_*(c*_*N*_*)}) < 0.83* or as TP otherwise. This filter targets SCS and chromosome fragments.
V. **Centromere width filter:** *W*_*cent*_*(c)* denotes the width of chromosome *c* at the top-ranked centromere candidate (see Fig. 3C). *c\** was classified as FP if *W*_*cent*_*(c*)/median({W*_*cent*_*(c*_*1*_*),…,W*_*cent*_*(c*_*N*_*)}) < 0.72* or as TP otherwise. This filter targets SCS and chromosome fragments.
VI. **Oblongness filter:** *S(c)* denotes the pair of side lengths of the minimum bounding rectangle enclosing the contour of chromosome *c* (see Fig. 3D). *c\** was classified as FP if 1 − *min(S(c*))/max(S(c*)) < 0.28* or as TP otherwise. This filter targets acrocentric chromosomes with SCS and some cases of overlapping chromosomes.
VII. **Contour symmetry filter:** Let *L(c)* denote the pair of arc lengths of contour halves produced by partitioning the contour of chromosome *c* at its centerline endpoints (see Fig. 3E). Classify *c\** as FP if *min(L(c*))/max(L(c*)) < 0.51* or as TP otherwise. This filter targets SCS.
VIII. **Intercandidate contour symmetry filter:** *L*_*C*_*(c)* denotes the pair of arc lengths of the contour regions of chromosome *c* that run between the traceline endpoints of its top 2 centromere candidates (see Fig. 3F). *c\** was classified as FP if *min(L*_*C*_*(c*))/max(L*_*C*_*(c*)) < 0.42* or as TP otherwise. This filter targets SCS and some cases of overlapping chromosomes.

**Figure 2.**
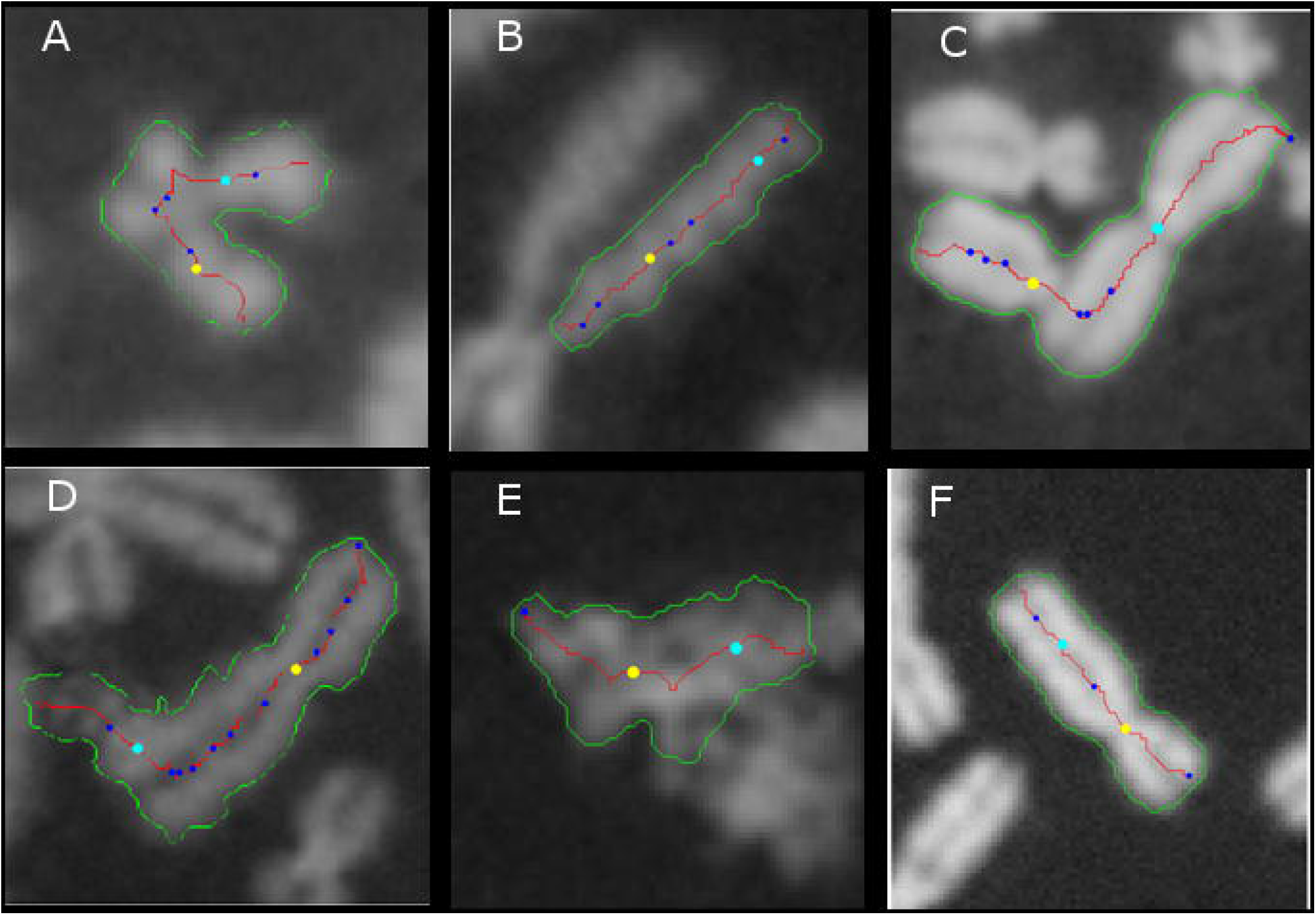
Examples of FPs in each morphological subclass. The subclasses are defined in the Methods 2. Chromosome contours are displayed in green, centerlines in red, top-ranked and 2^nd^-ranked centromere candidates in yellow and cyan, respectively, and other centromere candidates in blue. (A) SCS: An MC with SCS showing the characteristic localization of centerline along chromatid. (B) Chromosome fragment: Artifactual fragmentation of a chromosome caused by overaggressive image segmentation. (C) Chromosome overlap: Two touching MCs treated as a single DC (under-segmentation). (D) Noisy contour: The jagged contour due to poor image contrast is prone to introducing artifactual width constrictions. (E) Cellular debris: Incorrectly processed as a chromosome. (F) ML deficiency: An MC with no notable errors in contour or centerline.

**Figure 3.**
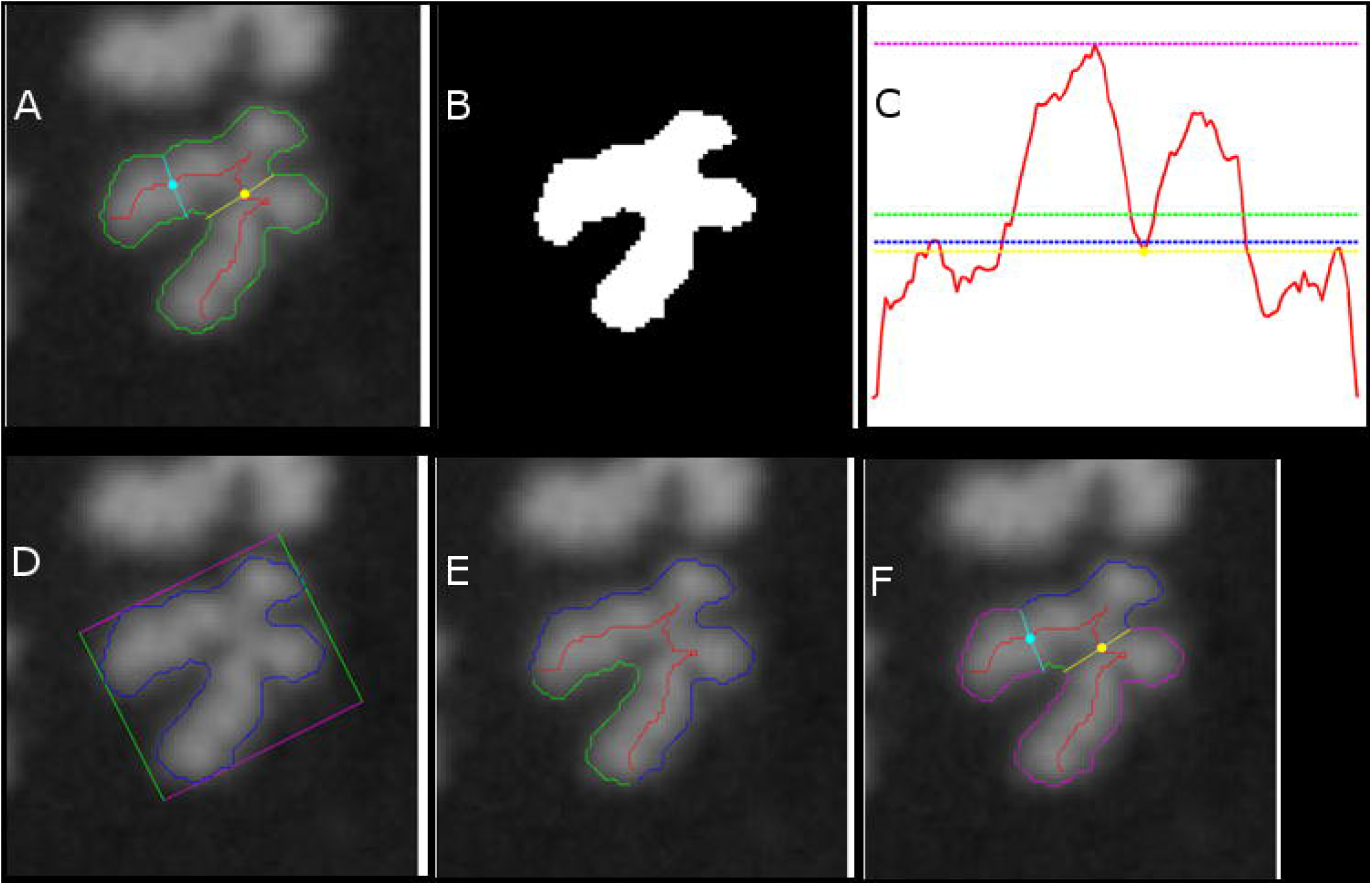
A visualization of DC filter scoresfor a particular FP. DC Filters are defined in Methods 3.1. (A) A processed FP (acrocentric chromosome with SCS), with contour in green, centerline in red, top-ranked centromere candidate and its width traceline in yellow, 2^nd^-ranked centromere candidate and its width traceline in cyan. (B) Filter I: Thresholded binary image of the chromosome is used to calculate pixel area (in white). (C) Filters II–V: Width profile along centerline is shown in red (horizontal axis plots centerline location, vertical axis plots width), with mean width in green (filter II), median width in blue (filter III), max width in magenta (filter IV), and width of top centromere candidate in yellow (filter V). (D) Filter VI: Contour in blue and its minimum bounding rectangle in magenta and green. (E) Filter VII: Partitioning of contour at centerline endpoints (intersection of red line with contour) into two segments, green and blue. (F) Filter VIII: Traceline endpoints of top 2 centromere candidates (intersection of yellow and cyan lines with contour) are used to partition contour into 4 segments (1 blue, 1 green, 2 magenta); relative arc lengths of blue and green segments are taken into consideration.

#### Incorporation into existing algorithms

After chromosome processing and MC-DC SVM classification^11^ but prior to dose determination, all DC chromosomes inferred by ADCI were analyzed with the proposed DC filters. DC filter scores exceeding TP thresholds were included in the dose determination, whereas DCs classified as FPs by any filters (inclusive “or”) were eliminated. DCs that were filtered out are outlined in yellow in the ADCI cell image viewer^11^ (Fig. 4).

**Figure 4.**
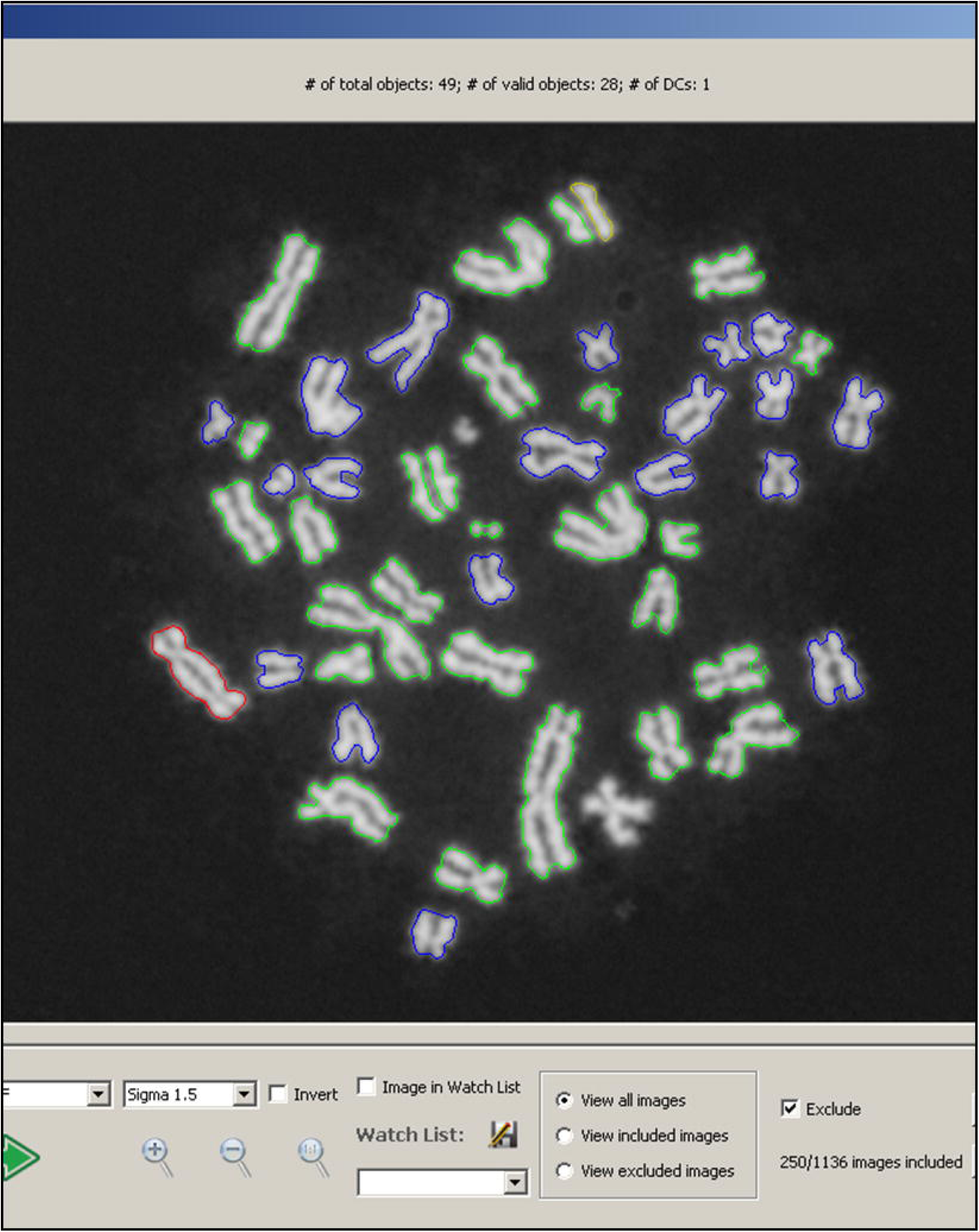
Cell image viewer in ADCI demonstrating example of a corrected FP DC. Graphical User Interface for viewing cell images within a sample processed by ADCI^11^. Valid segmented objects (generally chromosomes, but occasionally nuclei or debris) are shown with coloured contours. Red contours indicate predicted DCs, yellow contours indicate chromosomes that were initially classified as DC but removed by the FP filters (new), green contours indicate predicted MCs, and blue contours indicate objects that could not be further processed after segmentation. Beneath the image, new controls were added to allow manual inclusion/exclusion of images within a sample from dose analysis.

#### Determination of optimal filter subset

The proposed filters were not completely independent of each another, as some measures were related to the same chromosome segmentation features (i.e. width for filters II–V, contour symmetry for VII–VIII) and/or targeted the same morphological subclass (notably SCS). Thus, the “optimal” filter subset (termed “FP filters”) was defined as the subset of filters which maximized FP removal ability while minimizing redundant FPs. Performance for a given set of filters was the total percentage of FPs removed by any of its filters (inclusive “or”) in the *HC-mixed* image set (see Table 1). Using a forward selection approach, individual filters were added iteratively to identify those which produced the largest improvement in performance.

#### Evaluation of FP specificity on HC test samples

All objects removed by the FP filters in each image in HC samples INTC03S01, INTC03S08 and INTC03S10 (Table 3) were manually reviewed (Fig. 4). Filtered TPs and filtered objects with ambiguous classifications (TP or FP) were reviewed with another expert before final classification. For each sample, the number of filtered FPs was determined by subtracting number of filtered TPs from the total filtered count, and FP specificity was defined as the ratio of count of FPs to that of all filtered objects.

### 4) Dose estimation analysis

In ADCI, a pre-computed dose-response calibration curve is also used to estimate radiation absorbed in samples with unknown exposures^11^. For a given sample, ADCI calculates the mean response from total number of detected DCs divided by the number of cell containing images. Calibration curves can be generated from a set of calibration samples either by processing and calculating a response for each sample, or allowing the user to input the corresponding response, and fitting the dose-response paired data to a linear-quadratic curve by regression. Because sample preparation protocols vary between laboratories, dose estimation of test samples were performed with calibration curves generated by the same source^11^.

Distinct calibration curves were generated for each laboratory, either enabling or disabling FP filters, for the 0, 0.5, 1, 2, 3 and 4Gy calibration samples (see Table 2). Radiation doses of images obtained by HC for test samples (Table 3) were estimated using the HC calibration curve derived by ADCI after applying the same FP filters. A similar analysis was carried out for the 5 CNL test samples using the CNL calibration curve data.

### 5) Effect of filtering on manually image selected HC data

To investigate the impact of manual image selection on dose accuracy, we compared HC calibration curves derived from manually curated samples with the FP filters either enabled or disabled (Table 2). Manual curation of the HC samples was similar to manual image selection performed by CNL. Images were selected requiring: I) Complete complement of approximately 46 chromosomes, >40 segmented objects, <5 segmented objects from different nuclei if multiple nuclei present; II) Exclusion of “harlequin” chromosomes. Cells with unevenly stained sister chromatids cultured in the presence of bromodeoxyuridine (BrdU), which is indicative of 2^nd^ division metaphases, were excluded^10^; III) Well-spread, sharply-contrasted chromosomes with minimal sister chromatid dissociation. Only images with <5 incorrectly-segmented chromosomes were included, where incorrect segmentation was defined as chromosome overlaps (indicating poor spread), fragments (indicating sister chromatid dissociation) and overly-noisy contours (indicating poor image contrast); IV) Adequate chromatid condensation. Depending on the stage of metaphase arrest, the degree of chromosome condensation can differ^13,14^. Prometaphase cells have longer chromosomes, are less rigid, exhibit greater overlap and less well-defined centromere constrictions, all of which pose a significant challenges for automated chromosome classifiers^14,15^. Metaphase images with longer, thinner chromosomes (roughly corresponding to >500-band level^14^) were also excluded. Guidelines I-III and a minimum sample size of 500 cells were adopted from IAEA recommendations^12^, whereas guideline IV was added after preliminary inspection of HC calibration samples. Manual curation was performed within ADCI by retrospectively excluding images in processed samples from dose analysis (Fig. 4). For each sample, consecutive images meeting all criteria were evaluated until 500 images were accrued. DC classifications were hidden during image selection to minimize bias. After generation of the curated HC calibration curves, the radiation doses of the three HC test samples (Table 3) were re-estimated on the new curves, with and without the FP filters enabled.

### 6) Automating removal of suboptimal images by morphology filtering

Reference biodosimetry laboratories screen for interpretable metaphase cell images prior to DC analysis. Manual selection of images assures consistency and reliability of metaphase data, which increases analytic accuracy. As automated DC analysis can also be affected by variable cell image quality, excluding undesirable images in a sample would be expected to reduce FPs, and expected to more accurately estimate radiation exposures.

Image segmentation filters used empirically determined criteria to eliminate metaphase cells with characteristics that increased FP DCs. Image-level segmentation filters that threshold features I and II (below) were used to detect cells in prometaphase (relatively long and thin chromosome morphology), prominent sister chromosome dissociation, and highly bent and twisted chromosomes; another filter (III) detected overly-smooth contours characterized by images containing intact nuclei and otherwise incomplete chromosome sets. The total object count (IV) and segmented object count filters (V) fulfill general criteria for nearly normal metaphase images of approximately 46 chromosomes. These filters are used to exclude images with extreme object counts. Filter VI selects images based on effectiveness of chromosome recognition by ADCI.

Image level filters I-III are calculated in terms of their z-scores of all objects in an image. For any particular metaphase image *I\** in a sample containing *M* images, *{I*_*1*_*,…,I*_*M*_*}*, where *{c*_*1*_*,…,c*_*N*_*}* denote the set of *N* chromosomes within image *I\**. Additionally, *SD* denotes the standard deviation function, and *T* denotes the threshold SD common to all 3 filters that identifies outlier images. This SD value was set heuristically to 1.5 after by varying *T* after applying these filters to the HC2Gy calibration sample (Table 2). Similarly, suggested thresholds in filters IV-VI are also derived from experiences of testing multiple samples.

I. **Length-width ratio filter (LW)** defines the average length-width ratio of chromosomes in an image. For a given chromosome *c* in a given image *I* containing *N* chromosomes, *L(c,I)* denotes the arc length of the centerline of *c*, and *W*_*mean*_*(c,I)* denotes the mean value of the width profile of *c*. *MW(I) is defined as mean{L(c*_*1*_*,I)/W*_*mean*_*(c*_*1*_*,I),…,L(c*_*N*_*,I)/W*_*mean*_*(c*_*N*_*,I)}*. *I\** is removed if *MW(I*) > mean{MW(I*_*1*_*),…,MW(I*_*M*_*)} + T×SD{MW(I*_*1*_*),…,MW(I*_*M*_*)}*.
II. **Centromere candidate density filter (CD)** counts occurrences of centromere candidates in chromosomes. It eliminates images containing chromosomes with a high density of centromere candidates. For a given chromosome *c* in image *I* containing *N* chromosomes, *L(c,I)* denotes the arc length of the centerline of *c*, and *N*_*cent*_*(c,I)* denotes the number of centromere candidates along *c*. *CD(I)* is defined as the *mean{N*_*cent*_*(c*_*1*_*,I)/L(c*_*1*_*,I),…,N*_*cent*_*(c*_*N*_*,I)/L(c*_*N*_*,I)}*. *I\** is removed if *CD(I*) > mean{CD(I*_*1*_*),…,CD(I*_*M*_*)} + T×SD{CD(I*_*1*_*),…,CD(I*_*M*_*)}*.
III. **Contour finite difference filter (FD)** represents contour smoothness of chromosomes in an image. It eliminates images with prominent non-chromosomal objects with smooth contours, such as nuclei or micronuclei. For a given chromosome *c* in a given image *I* containing *N* chromosomes, *WP*_*D*_*(c,I)* denotes the set of first differences of the normalized width profile of *c* (range normalized to interval [0,1]). *WD(I)* is defined as the *mean{mean{abs{WP*_*D*_*(c*_*1*_*,I)}},…,mean{abs{WP*_*D*_*(c*_*N*_*,I)}}}*. *I\** is removed if *WD(I*) < mean{WD(I*_*1*_*),…,WD(I*_*M*_*)} – T×SD{WD(I*_*1*_*),…,WD(I*_*M*_*)}*.
IV. **Total object count (ObjCount) filter** defines the number of all objects detected in an image. Values lying outside of a threshold range are rejected to eliminate images with multiple metaphases or excessive cellular debris. Based on empirical analyses, the suggested object count range falls within the interval [40, 60].
V. **Segmented object count (SegObjCount) filter** defines the number of objects processed by GVF algorithm in an image. It is applied in the same way as filter IV. The suggested range for the object count interval is [35, 50].
VI. **Classified object ratio (ClassifiedRatio) filter** defines the ratio of objects recognized as chromosomes to the total number of segmented objects. It prevents images in which ADCI fails to process most chromosomes from being included. An image is removed if the value is less than a threshold of either 0.6 or 0.7, which is determined by the desired level of stringency for applicatoin of this filter.

#### Combining filters

Applying these filters sequentially to the same image distinguished the metaphase images for dose estimation from less optimal cells with increased FPs. This was done by combining the Z-scores of the image filters in a linear expression of features I-VI that provides an assessment of image quality. The resultant total score represents the degree to which a particular image deviates from the population of images in a sample:

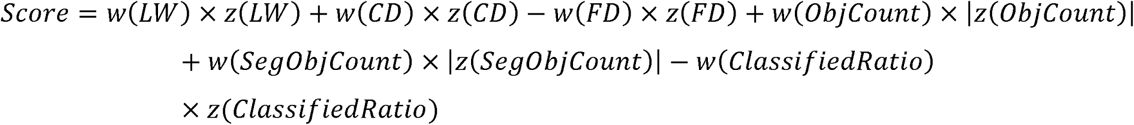

Each feature has a positive free parameter, weight, to adjust its contribution to the total score. The term LW determines that longer and thinner chromosomes in the image will increase the score, as do bending and twisted chromosomes due to the term CD. Lower chromosome concavity also drives the score higher because of FD term. Object count and segmented object count describe chromosome positioning, separated sister chromatid level, etc. Assuming the majority of images in a sample are good images, these terms will result in higher scores for images exhibiting either incomplete, multiple cells or severe sister chromatid separation. The last terms produce high scores for images that the algorithm does not process accurately. Images with smaller combined score are of higher quality. The weights used are identified by evaluating many possible weights and selecting those that minimize the error in curve calibration. The weights obtained are optimal for calibration samples, which will perform well on test samples, subject to the condition that the calibration and test samples have comparable chromosome morphologies. The score, however, cannot be used for inter-sample image quality comparisons, as z-scores are normalized within a sample.

Another, more general method was also developed to assess metaphase images separately from other images in the same sample. Image morphology is the primary consideration in assessing metaphase image quality. The most common problems in poor quality metaphase cells are severe sister chromatid separation, excessive chromosome overlap, fragments of chromosomes in image segmentation, and multiple cells or incomplete cells in the same image. They result in changes in either the number of objects or areas of objects. For instance, chromatid separation and chromosome fragments cause more objects to be present in an image while areas of some objects are smaller than normal. Chromosome-overlaps reduce the number of objects, but their areas exceed those of discrete chromosomes.

To derive this novel quality measure, we exploited the general property that the different chromosome lengths are approximately proportionate to the known base-pair counts of each complete human chromosome. By comparing the distribution of observed chromosome object lengths with the gold standard derived from the lengths obtained from the human genome sequence, we can assess the overall quality chromosome segmentation of each cell. This assumption sets aside chromosome abnormalities which result from radiation exposure, which will be distributed randomly among cells analyzed, because the cells are synchronized and harvested after a single division. The actual chromosome lengths are difficult to measure accurately in images, so instead, individual chromosomes are approximated according to their corresponding chromosome areas (in pixels). Therefore, the area of an object in a metaphase image is used as a surrogate for which chromosome it represents. Once noisy non-chromosomal objects, nuclei and large overlapped chromosome clusters are removed, areas of the remaining objects are then calculated based on their fractions to the total area of all chromosomes, as overlapping chromosomes and chromatid separation do not significantly affect the total area of objects in each metaphase image. We bin the chromosomes in metaphase cell into three categories corresponding to the known cytogenetic classification system^16^: group A and B (AB), group C (C) and groups D, E, F, and G (DG). A chromosome in category AB contains more than 2.9% (determined by the shortest B group chromosome) of total base-pairs in the complete chromosome set. A chromosome in category C has less than 2.9% (determined by the longest C group chromosome) but more than 2% (determined by the shortest C group chromosome) of total base-pairs in the set. Any chromosome in category DC contains fewer than 2% (determined by the longest D group chromosome) of the total base-pairs. These thresholds 2.9% and 2% are acceptable for the X and Y chromosomes, respectively. We apply these thresholds to object areas to count the number of chromosomes in each category in a metaphase image. An ideal metaphase image will have 10 AB chromosomes, 16 C chromosomes and 20 DG chromosomes if the individual is female, or 10 AB chromosomes, 15 C chromosomes and 21 DG chromosomes if the individual is male. Images with chromosome overlap will tend to have increased AB chromosome counts, while images with sister chromatid separation will likely have elevated DG chromosome counts. The morphological quality of a metaphase image can be measured by comparing its chromosome categorizing result to the female/male standard. In practice, we treat the categorizing result of an image as a 3-element vector and calculate the Euclidean distance to the standard. A larger distance corresponds to a less satisfactory image, and we find that this measurement is universal for metaphase images from different samples.

When images in a sample are sorted, by either combined z-score or by chromosome group bin area measurement, a certain number of top ranked images can then be selected for dicentric chromosome analysis. Complex image selection models can be created by filtering images first with filters and then selecting a certain number of top scoring images.

### 7) Sample Quality Confidence Measurement

Metaphase image artifacts such as sister chromatid separation and chromosome fragmentation interfere with the ability to correctly identify dicentric chromosomes, and compromises the reliability of dose estimates. This dependence of dose estimation accuracy on sample image quality motivates objective tests to evaluate and flag data from lower quality samples and exclude such images from analysis. Samples exposed to low LET whole-body irradiation, typically seen in radiation incidents, exhibit DCs frequencies that follow a standard Poisson distribution^17^ of DCs per cell. Deviation from the expected Poisson distribution can thus be attributed to failure to accurately recognize and account for DCs within the sample or by artifacts in the sample that are not eliminated by the software. Following this principle, we devised a sample quality evaluation method based on the conformity of the DC count frequency distribution in each sample to a theoretical Poisson distribution, as follows:

The number of DC occurrences in a cell is constructed as a probability model in a sample. It is a discrete statistical model as the number of events can only be integers. The appearance of any DC is assumed to be independent of other DCs that may form. The rate at which DCs occur is constant for a single sample at a given radiation dose for full-body irradiation. The model of DCs per cell detected by ADCI can therefore be approximated by a Poisson distribution. The Poisson λ parameter is obtained from the average number of DCs per cell in a sample.

The observed DC distribution detected by ADCI is compared with the Poisson distribution using the Pearson chi-squared goodness-of-fit test. The test indicates the probability of observing the observed data under the null hypothesis that they are Poisson distributed. Samples without at least 1 cell image having >1 DC cannot be analyzed, due to insufficient degrees of freedom. A smaller p-value means the hypothesis is less likely and that the DC detection results for that sample are less reliable. Very low p-values at or below α = 0.01 (99% confidence level) reject the null hypothesis and indicate low quality samples.

## RESULTS

### Application of chromosome morphology filters to remove FPs

False positive DCs (n=97) from a low dose set metaphase images were classified to uniquely identify, and ultimately eliminate these objects. Chromosomal morphological subclasses (Fig. 3) included those exhibiting excessive sister chromatid separation (I, n=51), fragmentation (II, n=10), overlap (III, n=17), noisy contours (IV, n=5), cellular debris (V, n=4), and inaccurate recognition by the centromere candidate^10^ and MC/DC^6^ machine learning algorithms (VI, n=11).

Segmentation filtering criteria were applied to these images. Scale-invariant filters were tested to determine thresholds that selectively removed subclasses I-III without eliminating any TPs. Of the 51 SCS cases, 35 involved short, acrocentric chromosomes. FPs were distinguished from TPs based on either their lower relative pixel area or width (filters I–V), substantially non-oblong footprint (filter VI), or substantial contour asymmetry across the centerline (filters VII and VIII). For filters I-V, normalization to median scores of other objects in the same image performed similarly to normalization to other measures of central tendency (e.g. z-score, mean, and mode after binning scores). FPs were eliminated for each morphological subclass (Table 4), with most of the segmentation filters acting on the targeted subclass, however, the effects of each filter were not exclusive to those subclasses (Methods 3).

**Table 4.** Comparison of FP subclass targeting between proposed DC filters.

| DC filter designation** | No. FP DCs removed filtering by morphological subclass* |  |  |  |  |  |
| --- | --- | --- | --- | --- | --- | --- |
|  | SCS<br>(n=51) | Fragment<br>(n=10) | Overlap<br>(n=17) | Noise<br>(n=5) | Debris<br>(n=4) | ML (n=11) |
| I: Area | 19 | 6 | 0 | 0 | 3 | 1 |
| II: Mean width | 14 | 6 | 0 | 0 | 2 | 0 |
| III: Median width | 12 | 5 | 0 | 0 | 3 | 0 |
| IV: Max width | 23 | 8 | 0 | 0 | 3 | 0 |
| V: Centromere width | 8 | 3 | 0 | 0 | 3 | 0 |
| VI: Oblongness | 31 | 1 | 2 | 0 | 0 | 0 |
| VII: Contour symmetry | 11 | 0 | 0 | 0 | 0 | 0 |
| VIII: Intercandidate<br>contour symmetry | 43 | 2 | 3 | 1 | 1 | 2 |

\*See Methods 2 for description of each subclass. Calculated from *HC-low* image set in Table 1.
\*\*See Methods 3.1 for description of each filter.

To evaluate individual filter performance, the percentage of FPs removed by each filter was calculated for the *HC-mixed* image set (Table 5). A two-sample Kolmogorov–Smirnov test (K–S) was also performed for each filter (α=0.05) on the same data, where one sample consisted of the filter scores of all TPs (n=183) and the other sample consisted of the scores of all FPs (n=158). All 8 filters rejected the null hypothesis (Table 5), suggesting that FPs can be discriminated from TPs using empirically-thresholded filter scoring. Application of the intercandidate contour symmetry filter (Methods 3.VIII) achieved the largest overall reduction of FPs (44.9%), and eliminated the most SCS-induced FPs (43 of 51). The *max width* filter (Methods 3. IV) yielded the next largest reduction in FPs (27.8%) and was the most efficient filter for detecting fragmentation-induced FPs (8 of 10).

**Table 5.** Comparison of FP discrimination ability between proposed DC filters.

| DC filter designation** | 2-sample K-S, TPs/FPs,<br>p-value* | FP removed (%)* |
| --- | --- | --- |
| I: Area | $2.2 \times 10^{-18}$ | 22.2 |
| II: Mean width | $9.2 \times 10^{-10}$ | 16.5 |
| III: Median width | $3.3 \times 10^{-9}$ | 14.6 |
| IV: Max width | $3.3 \times 10^{-8}$ | 27.8 |
| V: Centromere width | $8.8 \times 10^{-3}$ | 13.9 |
| VI: Oblongness | $1.1 \times 10^{-24}$ | 27.2 |
| VII: Contour symmetry | $1.2 \times 10^{-8}$ | 10.1 |
| VIII: Intercandidate contour symmetry | $4.0 \times 10^{-30}$ | 44.9 |

\*Calculated from *HC-mixed* image set from Table 1.
\*\*See Methods 3.1 for description of each filter.

**Table 6.**
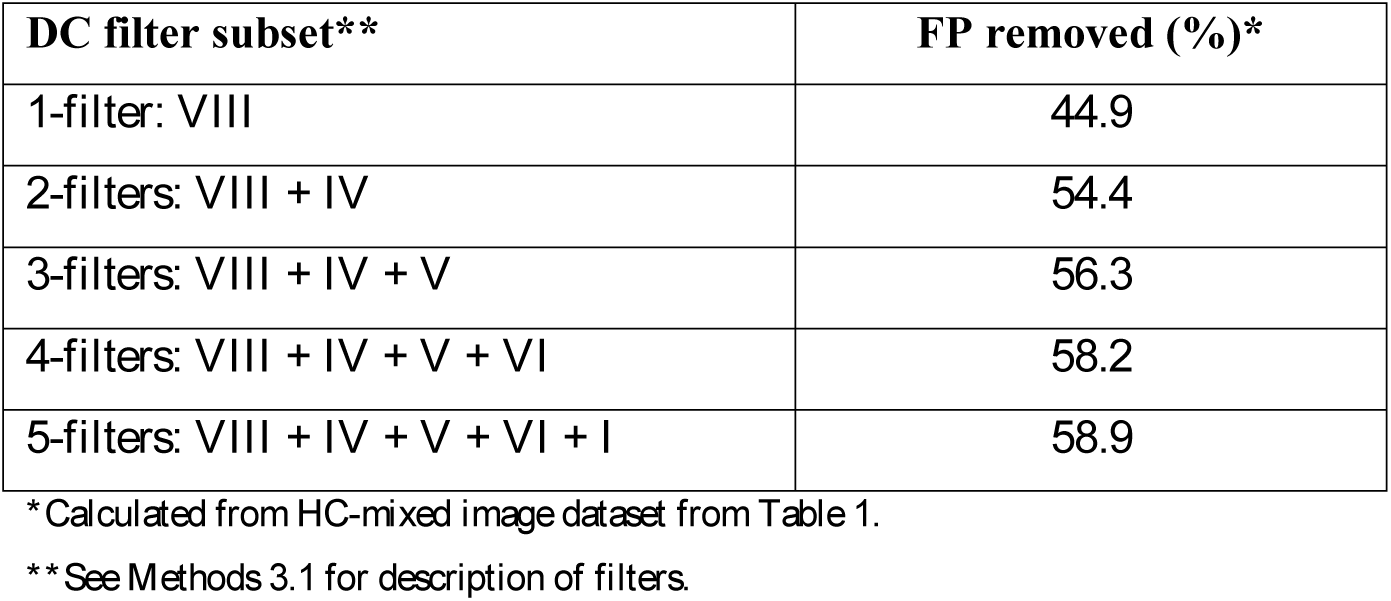
Forward selection results by combining subsets of DC filters.

| DC filter subset** | FP removed (%)* |
| --- | --- |
| 1-filter: VIII | 44.9 |
| 2-filters: VIII + IV | 54.4 |
| 3-filters: VIII + IV + V | 56.3 |
| 4-filters: VIII + IV + V + VI | 58.2 |
| 5-filters: VIII + IV + V + VI + I | 58.9 |

\*Calculated from *HC-mixed* image dataset from Table 1.
\*\*See Methods 3.1 for description of filters.

Additional FPs were eliminated by combining multiple segmentation filters (see Methods 3). Since individual filters were separately thresholded to avoid elimination of TPs (see Methods 3.2), the inclusive disjunction (logical “or” operation) of multiple filters had a negligible impact on TPs, while producing a stronger FP discriminator. Different combinations of filters were tested using forward selection. The best performing filter subset (collectively termed “FP filters”) consisted of a combination of 5 filters (I + IV + V + VI + VIII) that achieved a combined rate of FP removal of 58.9%. In comparison, the combination of filters IV + VIII accounted for most (54.4%) of the FPs eliminated, with incremental improvements resulting from ≤ 5 additional filters. Performance of these filters was evaluated on 3 sets of metaphase images (Table 7), consisting of 2 HC image sets (*HC-low* and *HC-high,* which were used during filter development) and an independent low dose image set from CNL. On average, 55 ± 9.6% of FPs were removed among all sets; individually the filters eliminated 52% of FPs from the CNL set, which was comparable to the HC sets (66% and 48% for low and high dose sets, respectively). All TPs were retained in each of the sets after processing of FPs (i.e. 100% specificity).

**Table 7.**
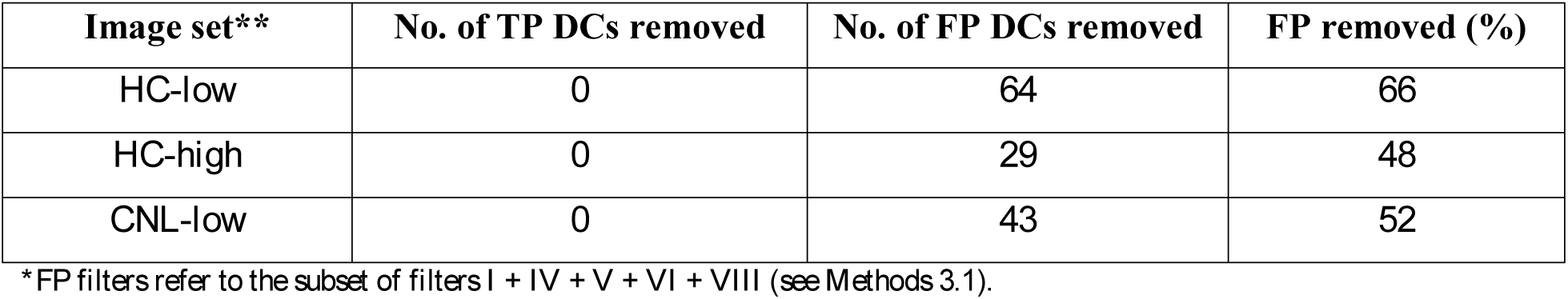
Performance evaluation of FP filters* on development and validation image datasets.

Dose-response calibration curves for HC and CNL data were generated in ADCI to investigate the effect of the filters on dose estimation accuracy (Fig. 5). Dose accuracy was assessed by determining the absolute error (absolute difference between dose estimate and true physical dose). For comparison, the dose estimates of 6 test samples (3 from HC, 3 from CNL) were compared which were either unfiltered and in which combinatorial FP filters were applied (Table 8). In samples that were manually curated by CNL, accuracy was also improved >2-fold by applying the 5 combined FP filters (average error decreased from 0.43Gy to 0.18Gy).

**Table 8.** Dose estimation of test samples, with and without FP filters* enabled.

|  | **HC samples\*\*** |  |  | **CNL samples\*\*** |  |  |
| --- | --- | --- | --- | --- | --- | --- |
| Physical dose (Gy) | 3.1 | 2.3 | 1.4 | 1.8 | 2.8 | 3.4 |
| Estimate, unfiltered (Gy) | 3.95 | 1.70 | 0.40 | 1.40 | 2.45 | 2.95 |
| Estimate, FP filters (Gy) | 2.65 | 1.4 | 0.20 | 2.20 | 2.80 | 3.50 |

\*FP filters refer to the subset of filters I + IV + V + VI + VIII (see Methods 3.1). Calibration curve image data was not curated or filtered. HC samples were unselected (INTC03S01, INTC03S08, and INTC03S10). The CNL samples were previously manually curated (INTC03S04 [n=448], INTC03S05 [n=500], and INTC03S07 [n=385]).
\*\*See Table 3 for sample details.

**Figure 5.**
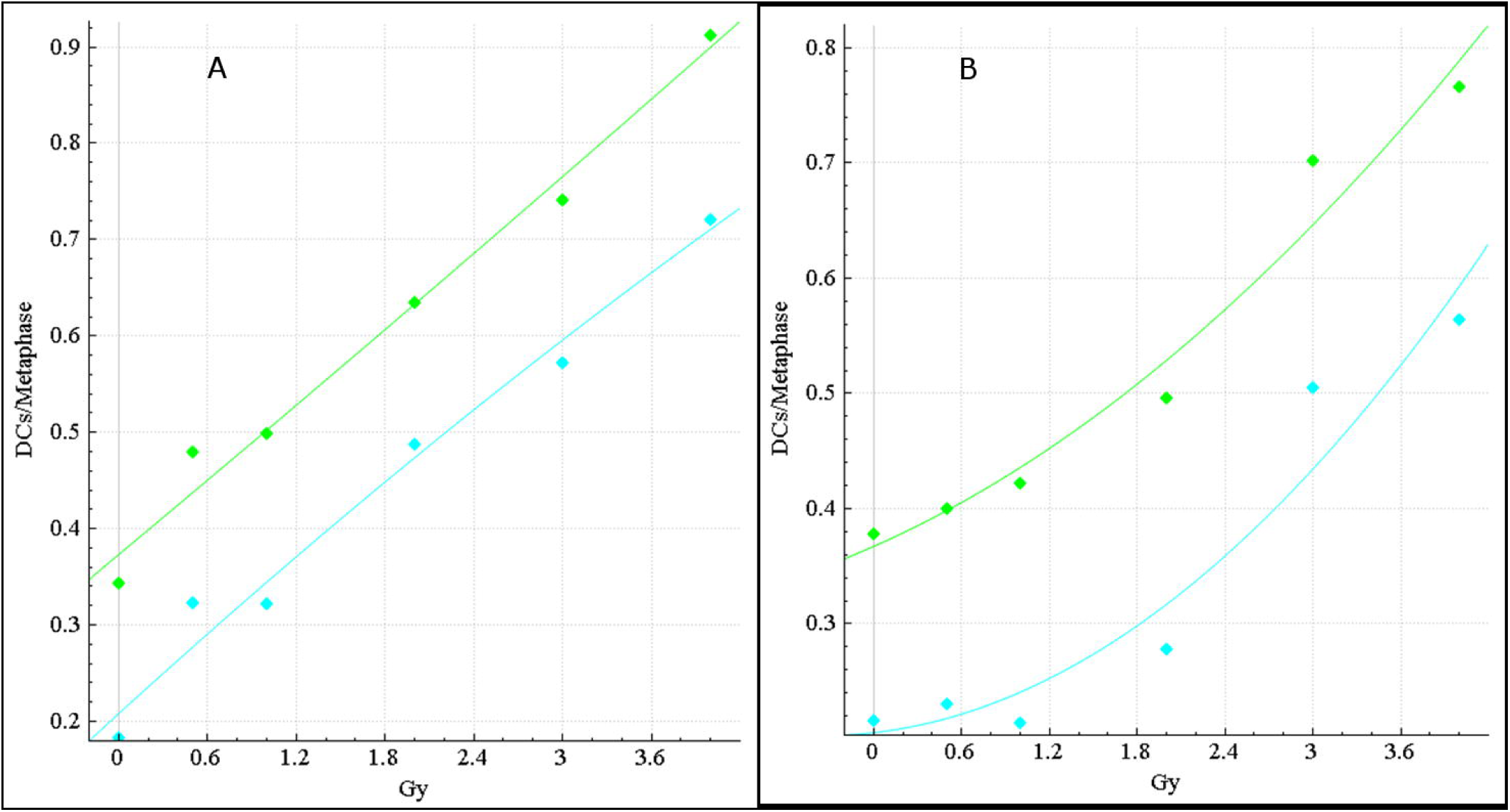
Calibration curves for HC and CNL samples. The dose-response calibration curves for (A) HC and (B) CNL metaphase cell image sample data. Response (mean DC frequency) on vertical axis, corresponding radiation dose (Gy) on horizontal axis. Green curves are based on unfiltered images, cyan curves were derived by recomputing DC frequencies after applying FP filters (filters I + IV + V + VI + VIII) to these datasets. HC curves are constructed by fitting a linear-quadratic curve through all HC calibration samples, CNL curves are similarly constructed from CNL calibration samples (refer to Table 2). The CNL curves consistently show a more pronounced quadratic component than the HC curves, which exhibit a nearly linear response. After applying FP filters (cyan), the curves show a diminished dose-response (green), due to elimination of some detected FP DCs.

The dose accuracy in the HC samples was impacted by addition of these filters (mean absolute error increased from 0.85Gy to 1.03Gy). One explanation was either the filters were removing many TPs inadvertently, or FPs removed by the filters were offsetting previously undetected DCs (false negatives) in the HC samples. All objects eliminated with these filters in the 3 HC samples were reviewed and classified as either TP or FP, and the FP specificity across the samples was determined (Table 9). Similar to earlier findings, the FP filters exhibited very high specificity for FPs (97.7–100%), indicating that the filters retained high specificity for TPs in the HC samples.

**Table 9.** Specificity of FP filters* in HC test samples.

| **Image sample\*\*** | **Total no. of chromosomes removed** | **No. of TPs removed** | **No. of FPs removed** | **Specificity for FPs** |
| --- | --- | --- | --- | --- |
| INTC03S01 | 193 | 0 | 193 | 100% |
| INTC03S08 | 133 | 3 | 130 | 97.7% |
| INTC03S10 | 143 | 2 | 141 | 98.6% |

\*FP filters refer to the subset of filters I + IV + V + VI + VIII (see Methods 3.1).
\*\*See Table 3 for sample details.

We hypothesized that the difference in image selection protocols was responsible for the discrepancies seen in classification performance and dose estimation accuracy between the two sources. While CNL manually selected for images deemed suitable for DCA analysis, image selection at HC was done with an automated metaphase classifier that effectively removed only images lacking metaphases (see Methods 1). Manual review of images in the HC and CNL samples confirmed noticeable differences in image quality: In concordance with findings from our previous study^1^, CNL data contained more images with well-spread, minimally-overlapping chromosomes, and fewer images with extreme SCS and chromosome fragments (complete dissociation of sister chromatids). The HC data contained a greater percentage of high-band-level (less condensed) chromosomes, characteristic of prometaphase/early-metaphase cell images. These chromosomes were the source of many unfiltered FPs, due to the lack of a strong primary constriction at the centromere.

A new set of HC calibration curves were then generated from manually curated, selected images from calibration samples (Fig. 6). Images were excluded based on IAEA criteria^17^, along with cells exhibiting long chromosomes in early prometaphase^16^ (Methods 5). (Table 10). Dose estimation accuracy of the HC samples (INTC03S01, INTC03S08 and INTC03S10) was significantly improved by enabling the 5 FP segmentation filters (mean unfiltered absolute error was 0.37Gy, and was 0.15Gy with the filters; Table 10). Therefore, application of FP filters to both CNL and curated HC data led to > 2-fold reduction in the mean absolute error of the estimated dose (*p* = 0.024, paired two tailed t-test).

**Table 10.** Dose estimates and deviations from physical dose for HC test samples after applying image selection models.

| **Image selection model** | **INTC03S01**<br>**3.1<sup>^</sup>** | **INTC03S08**<br>**2.3** | **INTC03S10**<br>**1.4** | **INTC03S04**<br>**1.8** | **INTC03S05**<br>**2.8** | **INTC03S07**<br>**3.4** |
| --- | --- | --- | --- | --- | --- | --- |
| All images | 2.65, -0.45 | 1.3, -1.0 | 0.2, -1.2 | 3.2, +1.4 | 2.1, -0.7 | 4.5, +1.1 |
| Combined z score, weight [5, 2, 4, 3, 4, 1], top 250 | 2.85, -0.25 | 1.8, -0.5 | 1.05, -0.35 | 4.65, +2.85 | 2.5, -0.3 | 5, +1.6, out of bounds |
| Combined z score, weight [4, 3, 4, 5, 2, 1], top 250 | 3.0, -0.1 | 1.8, -0.5 | 0.95, -0.45 | 4.95, +3.15 | 2.35, -0.45 | 5, +1.6, out of bounds |
| Combined z score, weight [1, 2, 1, 5, 1, 5], top 250 | 1.55, -1.55 | 1.35, -0.95 | 0.4, -1.0 | 4.4, +2.6 | 2.7, -0.1 | 5, +1.6, out of bounds |
| Chromosome group bin method, top 250 | 2.6, -0.5 | 2.3, +0.0 | 1.05, -0.35 | 1.6, -0.2 | 2.8, +0.0 | 2.2, -1.2 |
| Filters I-VI | 2.05, -1.05 | 0.95, -1.35 | 0.35, -0.95 | 0.6, -1.2 | 2.05, -0.75 | 1.15, -2.25 |
| Filters I-III & chromosome group bin method, top 250 | 3.0, -0.1 | 2.0, -0.3 | 0.95, -0.45 | 1.85, +0.05 | 2.95, +0.15 | 2.35, -1.05 |
| Manual selection | 2.85, -0.25 | 2.3, +0.0 | 1.05, -0.35 | 3.95, +2.15 | 2.45, -0.35 | 4.0, +0.6 |

<sup>^</sup> Sample identifier, physical dose (Gy). FP filters were enabled.

**Figure 6.**
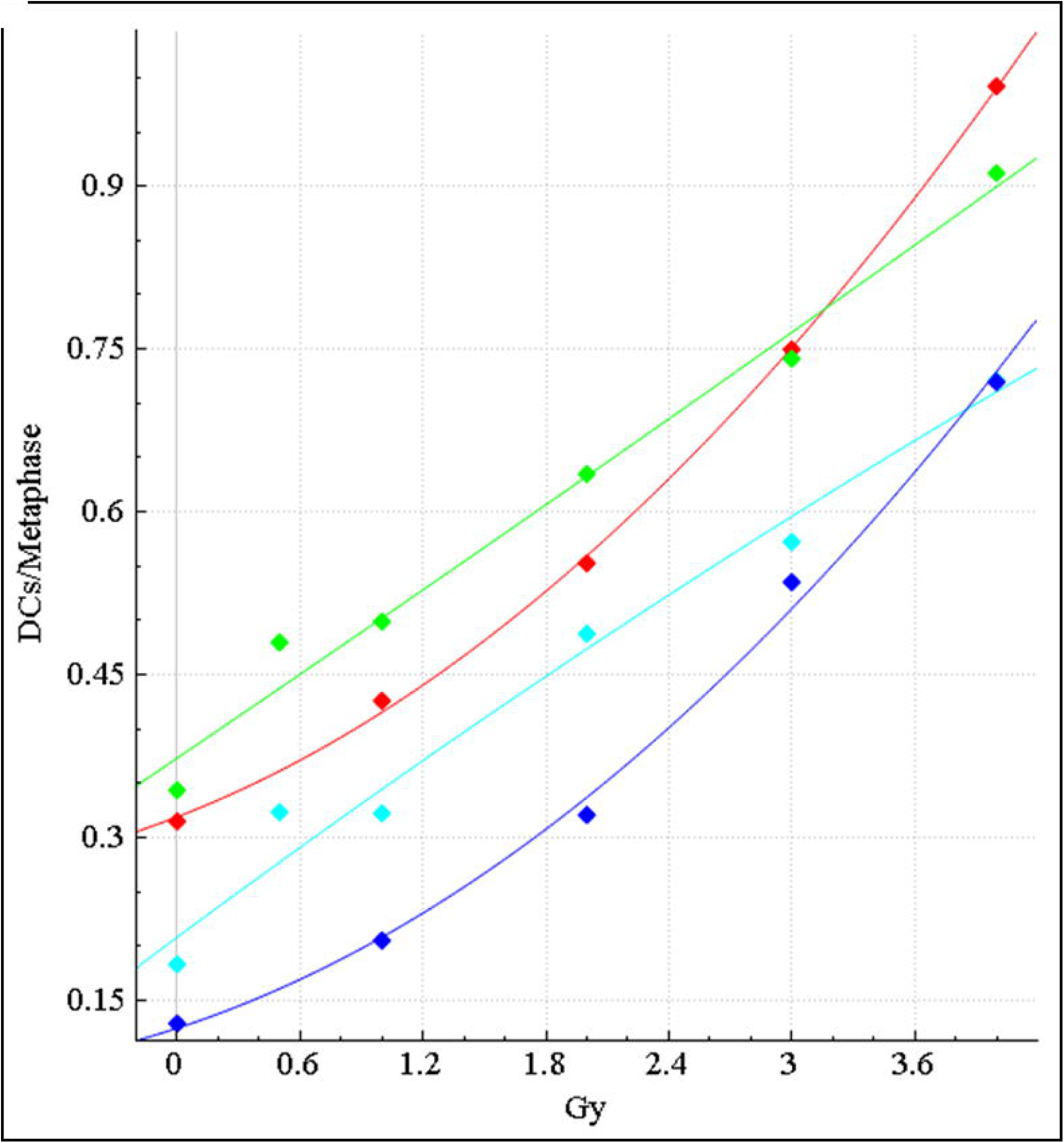
Original vs. manually curated calibration curves for HC samples. The dose-response calibration curves for HC sample data, with and without FP filters applied, before and after curation. Response (mean DC frequency) on vertical axis, corresponding radiation dose (Gy) on horizontal axis. Green curve is not curated and includes all images, cyan curve is not curated and applies FP DC filters, red curve is curated, but unfiltered, and blue curve is curated and FP filters have been applied. Uncurated curves were generated from 0, 0.5, 1, 2, 3 and 4Gy calibration image data (see Table 2). Curated curves were generated from the same data (however 0.5Gy was not included) after lower quality images were manually removed (see Methods 6). After manual curation, the curves show a stronger quadratic component, similar to the CNL curves (see Fig. 5).

### Application of Image Selection Models

Assessment of image selection was challenging, as no objective standard exists. Cell selection by cytogenetic experts is based on their knowledge of metaphase chromosome conformation, sensitivity, and even individual preferences in interpreting images which are sometimes inconsistent. Therefore, image selection methods were evaluated through dose estimation of filtered test samples and comparisons with known physical exposures. The images in all calibration and test samples were processed by the same image selection method. Dose estimates of test samples are calculated using a curve fit to calibration samples. Dose estimation errors indicate the accuracy of dicentric chromosome detection, and consequently imply the effectiveness of image selection method.

To rank images with the combined z-score method, a weight vector corresponding to each of the 6 filters comprising the total score was first determined. Optimal weights were obtained by searching a large number of possible values among the set of HC calibration samples for those exhibiting smallest residuals when fit to the curve. The potential weights were defined as integers ranging from [1, 5]. This limited the search space and eased computational complexity, but nevertheless ensured that diverse combinations of weights were evaluated. In experiments, three optimal weight vectors, namely [5, 2, 4, 3, 4, 1] [4, 3, 4, 5, 2, 1] and [1, 2, 1, 5, 1, 5], were used for dose estimation.

After images were assigned scores and sorted according to their combined z-scores (or by the chromosome group bin method-see below), the 250 top ranked images were subsequently selected to determine dicentric aberration frequency for that sample. An adequate number of top ranked images are selected to provide sufficient images to generate a reproducible DC frequency for that sample. The top ranked image set also has to effectively remove poor quality images that could distort the DC frequency. IAEA has recommended at least 100 DCs be detected for samples with physical doses >1 Gy. In practice, laboratories score >250 images, but often more. Considering the total number of images in a sample ranges from 500 to 1500, we found that selecting the 250 top scoring images gave satisfactory results. Figure 7 indicates that the DC frequency for the HC3Gy calibration sample stabilizes after at least 250 to 300 top images were included. Similar results were obtained for other test and calibration samples (not shown). DC frequencies can differ between image selection methods because each method can select different images. When the number of top ranked images significantly exceeds 300 images, differences between the specific image selection methods are minimized as they share increasing numbers of selected images. Unfiltered randomly sampled images from this sample tend to exhibit higher overall DC frequencies due to increased numbers of FP DCs.

**Figure 7.**
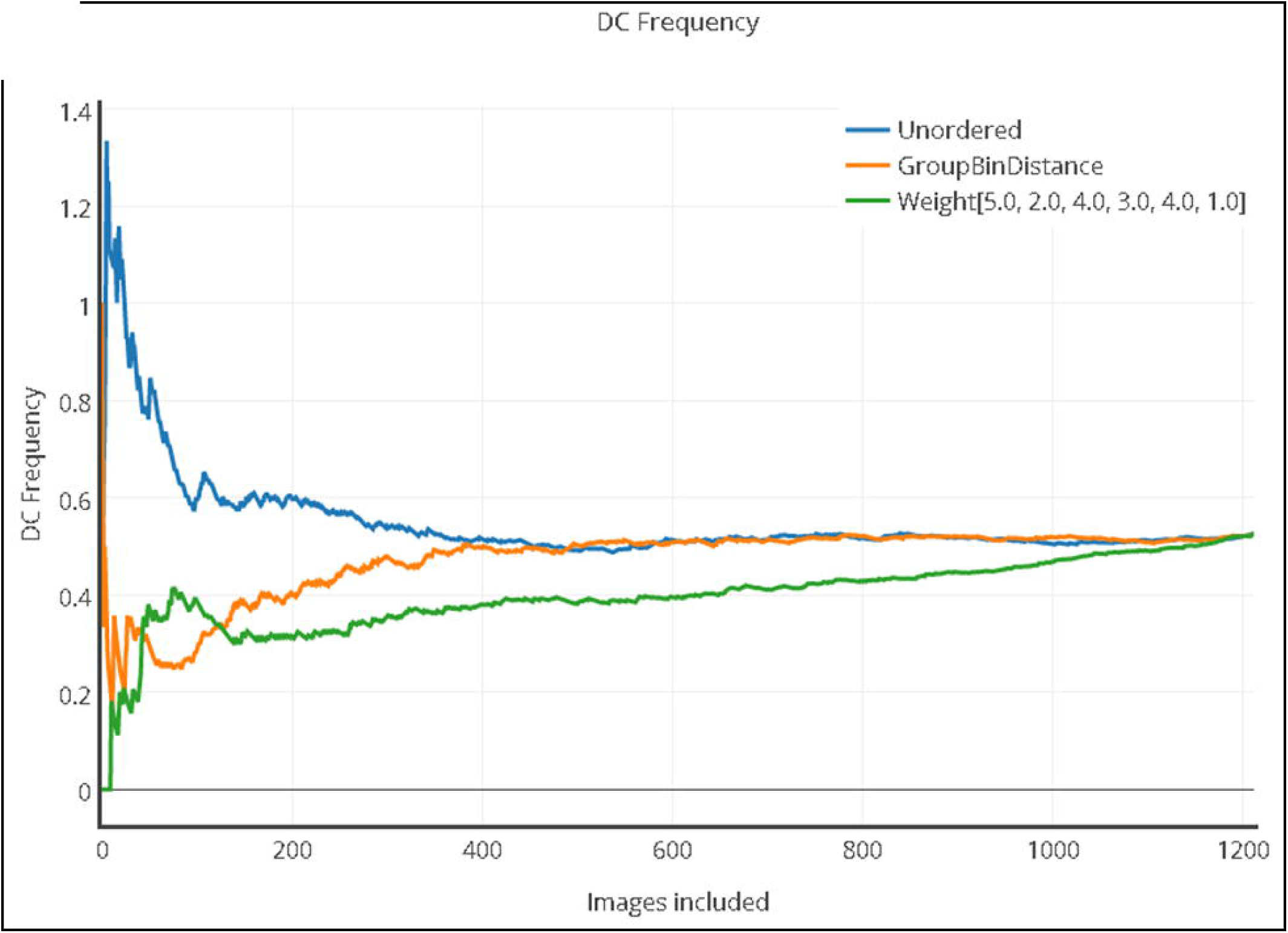
Relation between DC frequency (y-axis) and number of included top images (x-axis) when images are ranked by different scoring methods, in sample HC3Gy. Blue, orange and green curves correspond to unordered images (alphabetic order of image names), images sorted by group bin method and images sorted by combined z-score method, respectively. Figure was generated using <u>Plotly</u> software.

The deviations of estimated doses of all of the HC and CNL test samples, respectively, from physical doses, were determined for various ADCI image selection models (Tables 10 and 11). For comparison, the dose estimation results of unselected, comprehensive sets of images for each sample are presented. Deviations of ≤ 0.5 Gy from their calibrated physical dose are acceptable for triage biodosimetry^5,12^. For the unfiltered HC samples, the average absolute error is 0.8 Gy, with a single sample, INTC03S01, fulfilling the triage criteria. The image selection model that combines filters I-III and chromosome group bin method produces the best result. Dose estimates for four samples (INTC03S01, INTC03S08, INTC03S10 and INTC03S05) are acceptable. The combined z-score method with the filter weights: [1, 2, 1, 5, 1, 5] resulted in the least accurate estimates. Here, the average error is ∼1 Gy, and only INTC03S05 had an acceptable dose estimate. Of the five unfiltered CNL samples, only INTC03S08 had an acceptable dose estimate. After applying image selection models, a pan-filter set using all of the available filters I-VI gave the most accurate results. The average absolute error was ∼0.3 Gy, and 4 of 5 samples (INTC03S08, INTC03S04, INTC03S05 and INTC03S07) exhibited doses in the acceptable range.

**Table 11.** Dose estimates and deviations from physical dose for CNL test samples after applying image selection models.

| **Image selection model** | **INTC03S01**<br>**3.1<sup>^</sup>** | **INTC03S08**<br>**2.3** | **INTC03S04**<br>**1.8** | **INTC03S05**<br>**2.8** | **INTC03S07**<br>**3.4** |
| --- | --- | --- | --- | --- | --- |
| All images | 4, +0.9, out of bounds | 2.6, +0.3 | 2.45, +0.65 | 3.6, +0.8 | 4, +0.6, out of bounds |
| Combined z score, weight [5, 2, 4, 3, 4, 1], top 250 | 3.9, +0.8 | 2.8, +0.5 | 2.1, +0.3 | 3.05, +0.25 | 3.55, +0.15 |
| Combined z score, weight [4, 3, 4, 5, 2, 1], top 250 | 4, +0.9, out of bounds | 2.75, +0.45 | 1.7, -0.1 | 3.05, +0.25 | 3.9, +0.5 |
| Combined z score, weight [1, 2, 1, 5, 1, 5], top 250 | 3.6, +0.5 | 2.4, +0.1 | 0.65, -1.15 | 2.35, -0.45 | 3.05, -0.35 |
| Chromosome group bin method, top 250 | 4, +0.9, out of bounds | 2.8, +0.5 | 1.8, +0.0 | 2.55, -0.25 | 4, +0.6 |
| Filters I-VI | 3.75, +0.65 | 2.8, +0.5 | 1.95, +0.15 | 3.05, +0.25 | 3.4, +0.0 |
| Filters I-III & chromosome group bin method, top 250 | 4, +0.9, out of bounds | 2.75, +0.45 | 1.65, -0.15 | 2.25, -0.55 | 3.95, +0.55 |
| Manual image curation | 3.7, +0.6 | 2.85, +0.55 | 2.2, +0.4 | 2.8, +0.0 | 3.5, +0.1 |

<sup>^</sup> Sample identifier, physical dose (Gy). FP filters were enabled.

Image selection rejects poor images and reduces FP DCs if sufficient quantities of images remain to provide reliable DC frequencies. Although >250 images were usually present after scoring and ranking, application of image filters can result in fewer remaining images for analysis. After applying the pan-filter set, sample CNL-INTC03S08 consisted of 195 metaphase cells. After applying the combined image selection model to the HC samples, sample HC-INTC03S07 consisted of only 109 metaphase cells. This sample was relatively lower quality than others in this set, and the unfiltered set of metaphase images was smaller than the recommended minimum, consisting of 477 cells (Table 12).

**Table 12.** Goodness of fit Poisson scores* of unfiltered, manually- and ADCI-filtered image sets for calibration and test samples.

| Sample | Unfiltered images | Manual image selection | Automated image selection |
| --- | --- | --- | --- |
| HC0Gy | 1.333e-15 | 2.240e-01 | No DCs detected |
| HC05Gy | 4.084e-32 | unavailable | 1.146e-02<sup>^</sup> |
| HC1Gy | 1.669e-18 | 9.996e-01 | 1.049e-01<sup>^</sup> |
| HC2Gy | 4.019e-64 | 2.072e-01 | 7.618e-04<sup>^</sup> |
| HC3Gy | 2.873e-02 | 4.642e-01 | 6.112e-01<sup>^</sup> |
| HC4Gy | 2.596e-04 | 2.215e-01 | 3.127e-01<sup>^</sup> |
| HC-INTC03S01 | <2.225e-308<sup>+</sup> | 9.052e-01 | 1.170e-01<sup>^</sup> |
| HC-INTC03S08 | 1.236e-01 | 4.573e-01 | 8.153e-01<sup>^</sup> |
| HC-INTC03S10 | 8.873e-01 | 3.895e-01 | 2.113e-01<sup>^</sup> |
| HC-INTC03S04 | <2.225e-308 | <2.225e-308 | 5.533e-01<sup>^</sup> |
| HC-INTC03S05 | 1.103e-06 | 7.262e-05 | 3.544e-03<sup>^</sup> |
| HC-INTC03S07 | <2.225e-308 | <2.225e-308 | 1.996e-04<sup>^</sup> |
| CNL0Gy | 5.174e-03 | 1.254e-01 | 3.071e-01<sup>#</sup> |
| CNL05Gy | 1.656e-157 | 1.236e-01 | 5.955e-32<sup>#</sup> |
| CNL1Gy | 9.801e-30 | 1.496e-03 | 1.597e-06<sup>#</sup> |
| CNL2Gy | 2.340e-147 | <2.225e-308 | 4.488e-02<sup>#</sup> |
| CNL3Gy | 8.489e-07 | 6.820e-03 | 9.914e-01<sup>#</sup> |
| CNL4Gy | 5.151e-22 | 3.303e-02 | 1.826e-01<sup>#</sup> |
| CNL-INTC03S04 | 1.728e-60 | 1.933e-02 | 5.446e-02<sup>#</sup> |
| CNL-INTC03S05 | 2.743e-09 | 5.243e-02 | 3.253e-01<sup>#</sup> |
| CNL-INTC03S07 | 6.671e-10 | 4.248e-05 | 4.725e-01<sup>#</sup> |
| CNL-INTC03S01 | <2.225e-308 | 3.735e-03 | 7.627e-11<sup>#</sup> |
| CNL-INTC03S08 | 5.253e-16 | 7.678e-01 | 7.768e-01<sup>#</sup> |

\* Poisson score is the p-value of chi-square goodness of fit (without merging bins) of observed distribution of DCs/cell vs.
Poisson distribution determined from average DC frequency. Filtering parameters chosen for each laboratory exhibit dose estimates that are closest to the physical dose: <sup>^</sup>HC image sets were filtered with morphological filters I-III and by chromosome group bin score; <sup>#</sup>CNL image sets were filtered with morphological filters I-VI. <sup>+</sup>Minimum positive floating value in Windows operating system.

### Sample Quality Assessment after Image Selection

To evaluate whether the image selection models improved sample quality, a Chi squared goodness of fit test was performed on the observed DC/cell vs. Poisson distributions for the CNL and HC samples, both prior to and after automated and manual image selection (Table 12). Manual image selection for CNL samples was performed by CNL during sample preparation, while image selection for HC samples was performed on unselected datasets (see Methods 5; samples HC-INTC03S01, HC-INTC03S08, HC-INTC03S10 were analyzed, despite <500 images being available). For each laboratory, the best performing image selection models were used for FP and image level filtering (Tables 10 and 11). Image selection with filters I-III and chromosome group bin method was applied to HC sample data, whereas filters I-VI were applied to the CNL data. At the 1% significance level (i.e. Poisson goodness-of-fit, p ≤ 0.01), 86% (19 of 22) of unfiltered samples are significantly differed from the Poisson distribution, and 76% (13 of 17) of manually- and 77% (17 of 22) of automatically-selected samples samples did not differ; manually curated and uncurated sample groups also significantly differed from each other (p = 0.0021; one-tailed Wilcoxon Signed-Rank Test, α=0.05, n=17). Therefore, the Poisson goodness of fit measures changes in overall sample quality from image model selection. While the Poisson score is improved for all of the automatically selected datasets, the lowest quality samples (CNL1Gy, CNL05Gy, CNL-INTC03S01, HC-INTC03S05, HC-INTC03S07) were still rejected as Poisson-distributed after automated filtering.

## DISCUSSION

Automated biodosimetric methods aimed at detecting DCs can produce incorrect assignments because the algorithms cannot capture the full range of morphological variability inherent in chromosome images of metaphase cells. Accuracy of radiation exposure estimates using automated biodosimetry can be improved by image segmentation and filtering methods that remove suboptimal metaphase cell images and eliminate false positive DCs. This study implements and tests a set of morphology-based filters to eliminate FP DCs and unsuitable metaphase images for automated biodosimetry. Compared to results generated by the previous version of ADCI^11^, inclusion of these filters reduced FP DC rates by ∼55% across a wide range of radiation exposure levels. Additionally, we showed that these filters were highly specific for FPs in test image sets as well as actual patient samples (97.7–100%, n=6). Overall, the FP filters substantially improve DC classification accuracy.

This is because proposed segmentation filters successfully target SCS and chromosome fragments. In particular, the *intercandidate contour symmetry* filter is a very promising SCS detector, individually eliminating 84% of all SCS-induced FPs in our test dataset. It was noted that acrocentric chromosomes were disproportionally susceptible to SCS-induced errors compared to other chromosome types (69% of SCS cases despite making up only 22% of human chromosomes). Given the rarity of acrocentric TP DCs (due to width profile inaccuracies at the extreme ends of chromosomes^7–9^), filters targeting acrocentric or small chromosomes, in general (such as filters I and VI), can also be useful for reducing SCS-induced FPs.

Certain FP subclasses were commonly targeted by multiple filters. Redundancy among the segmentation features resulted in only subset of the filters being required to maximize elimination of FPs. Notably, filters II–V eliminated FPs based different definitions of chromosome width. The final combination of FP filters consisted of only 5 of the 8 originally proposed; however, it should be noted that a combination of only the *intercandidate contour symmetry* and *max width* filters achieved nearly the same level of FP detection in the test sample dataset, with the other filters having incremental benefit.

Scale-invariance is an obligate property for any object-level filter, since chromosome structures may vary between cells, individuals, and laboratory preparations. Scale invariance is also necessary to control for pixel-based chromosome measurements affected by condensation differences over the course of metaphase and differences in optical magnification. This principle was achieved by either using filter scores normalized to the median “raw” score of all objects within the same cell image (i.e. filters I–V), or in which scores were derived from ratios of two pixel-based measurements (i.e. filters VI–VIII).

Limitations of the current set of filters were revealed by differences in accuracy between the manually and automatically-selected images for dose estimation. For the previously manually curated CNL and HC samples, the FP object filters respectively reduced the average dose estimation error from 0.4Gy to <0.2Gy (with a maximum error of 0.4Gy). This placed the accuracy our software comfortably within the ±0.5Gy requirement for triage purposes^17^. However, applying the FP object filters alone to unselected HC metaphase data did not improve accuracy (average error increased by 0.15Gy). Thus, FP object filters alone did correct for inaccurate dose response estimates in all cases.

Variable cell image quality in some samples contributed to this source of error. Some unselected HC samples contained images with high levels of SCS, which upon processing produced large numbers incorrectly classified chromosome fragments. Image level filters I–V targeted these fragments, however they were not excluded based on their threshold values, because they comprised the predominant morphology within these particular cells. For similar reasons, object-level filtering was not suitable for elimination for removal of prometaphase images containing high resolution chromosomes (>800 band level). These observations suggested the need for image-level filters to select low quality images for removal in addition to the object-level filters.

Image quality is critical to the accurate DC detection. Manual inspection and quality control is common practice in cytogenetics and biodosimetry laboratories, but it is labor-intensive. Image-level filtering was automated to address this problem. These methods apply statistical thresholds to morphological features of chromosomes and non-chromosomal objects throughout a metaphase cell image. Image scoring methods select a defined number of top-ranked, processed images for dose estimation. The combined z-score method is a weighted sum of standard deviations below or above the mean score of objects in an image for each of the filter, and indicates relative image quality. The chromosome group bin method is a more general criterion that is calibrated to relative chromosome lengths (and area) in base pairs. ADCI evaluates the morphological deviation of chromosome area and ranks cell images relative to that expected from the standard, normal karyotype. These FP filtering and image scoring methods, which are referred to collectively as image selection models, can be applied either individually or in combinations within ADCI.

Significant improvement in accuracy of DC frequency is attributable to both FP elimination and image selection. Dose estimation errors with suitable image selection models in test samples consisting of at least 250 images are considerably reduced. The estimates are within the +/- 0.5 Gy window of the corresponding physical doses for the majority of samples we tested. The current models in ADCI generally provide reliable image quality control without manual intervention.

Automated image selection aims to simulate manual image curation. Experiments demonstrated that the proposed methods successfully improve dose estimates in test samples. At this point, automation does not quite achieve the same overall accuracy, especially for samples of variable quality. The respective differences in dose estimates, especially at exposures >2 Gy, are not significant. Automating image selection, nevertheless, offers unique advantages over manual image selection in terms of analytic uniformity and speed.

## ACKNOWLEDGEMENTS

This study was supported by the Build in Canada Innovation Program (Contract No. EN579-172270/001/SC) and CytoGnomix. The previous version of ADCI and development of algorithms were supported by the Western Innovation Fund; the Natural Sciences and Engineering Research Council of Canada (NSERC Discovery Grant 371758-2009); US Public Health Service (DART-DOSE CMCR, 5U01AI091173-0); the Canadian Foundation for Innovation; Canada Research Chairs, and CytoGnomix Inc.

